# NRGSuite-Qt: A PyMOL plugin for high-throughput virtual screening, molecular docking, normal-mode analysis, the study of molecular interactions and the detection of binding-site similarities

**DOI:** 10.1101/2025.01.23.634566

**Authors:** Gabriel T. Galdino, Thomas DesCôteaux, Natalia Teruel, Rafael Najmanovich

**Affiliations:** Department of Pharmacology and Physiology, Université de Montréal, Montreal, Canada

**Author notes:** These authors contributed equally (Gabriel T. Galdino, Thomas DesCôteaux).

## Abstract

**Summary:** We introduce NRGSuite-Qt, a PyMOL plugin that provides a comprehensive toolkit for protein modeling, virtual screening, normal mode analysis, and binding-site similarity calculations. Building on the original NRGSuite plugin for FlexAID, this updated version integrates five new functionalities: protein-protein and protein-ligand interaction analysis using Surfaces, ultra-massive virtual screening with NRGRank, binding-site similarity detection with IsoMIF, normal mode analysis using NRGTEN, and mutational studies through integration with the Modeller Suite. By merging these advanced tools into a cohesive platform, NRGSuite-Qt streamlines complex workflows and facilitates high-throughput computational studies within a single interface. Additionally, we benchmark a newer version of the Elastic Network Contact Model for normal mode analysis method ENCoM, utilizing the same 40 atom-type pairwise interaction matrix that is used in all other software. This version outperforms the default model in multiple benchmarking tests.

**Availability:** The Installation guide and tutorial is available at https://nrg-qt.readthedocs.io/

## 1 INTRODUCTION

Computational methods for evaluating protein structures and complexes are fundamental to drug discovery and understanding the mechanisms of various biological systems. However, analyzing all interactions and dynamics of a protein using traditional all-atom models can be extremely costly, depending on the size of the system studied and the amplitude of the movements of interest [1]. Recent advances in protein structure prediction and the wide availability of purchasable synthetic compounds have increased the demand for fast and reliable tools capable of modeling and comparing millions (or even billions) of systems at reduced computational costs [2, 3]. Coarse-grained (CG) methods offer an efficient alternative to traditional all-atom models. These methods use simplified representations of systems, reducing complexity and enabling the study of large biological systems in a shorter time.

Among these methods, docking software (DS) has emerged as a valuable tool for drug discovery. This approach uses scoring functions to enrich true binders in ligand libraries for targets with known structures [4]. The Flexible Artificial Intelligence Docking (FlexAID) software [5], previously introduced by our group, performs the docking of flexible ligands while also allowing flexibility in the target’s side chains. It has been extensively validated in various scenarios [6, 7] and was previously available in the original NRGSuite as a PyMOL plug-in for real-time docking [8, 9]. NRGSuite-Qt reintroduces an intuitive, user-friendly, and interactive interface for performing docking simulations for single ligands employing FlexAID. It also contains GetCleft [5], a software for cavity detection and binding site definition based on the SurfNet algorithm [10], providing seamless integration between cavity determination and the exploration of these cavities for small-molecule docking.

One recent FlexAID application was the identification of ligands targeting the Ebola VP35 protein in a virtual drug repurposing screening campaign. In this pipeline, ligands were first screened using FlexAID in the Chemical Component Dictionary (CCD) [11], followed by selection based on binding-site similarity analysis with IsoMIF [12], a software for binding-site similarity calculation based on the detection of binding site molecular interaction field similarities — also integrated in NRGSuite-Qt. Two molecules were identified to be capable of preventing the interaction between VP35 and ubiquitin, reducing viral replication both in vitro and in vivo [13]. Another noteworthy application was in the identification of Toyocamycin as a selective CDK9 inhibitor in cancer cells, with a binding site characterization that opens the possibility for the design of novel CDK9 inhibitors [7].

The FlexAID scoring function (FSC) relies on a pairwise interaction matrix comprising 40 SYBYL-based atom types. This interaction matrix was derived through Monte Carlo optimization based on known ligand-protein crystal structures available in the Protein Data Bank (PDB) and shows a strong correlation with established chemical data [5]. The FSC was implemented in Surfaces [14], a CG method for analyzing protein-protein (PPI) and ligand-protein (LPI) interactions. Surfaces has demonstrated performance comparable or superior to molecular dynamics-based methods (e.g., MM/PB-SA, MM/GB-SA, FEP+ and gRINN analysis) in predicting the effects of mutations on PPI and binding affinity, particularly in evaluating the impact of mutations on interactions involving the SARS-CoV-2 Spike protein. This software was also successfully used to identify mutations that disrupted interactions between ligands and VP35 without affecting its interaction with ubiquitin [13].

NRGRank [15] uses a simplified version of FSC. It is a structurally-informed ultra-massive virtual screening software that can screen large ligand libraries at extremely low computational cost resulting in a ranked list of ligands and approximate poses (actual binding poses can be obtained subsequently with FlexAID). This software was validated in different scenarios and is capable of screening 50 000 ligands per hour on a common laptop (8 cores).

More recently we employed the FSC within Elastic Network Contact Model (ENCoM) [16] as part of the the Najmanovich Research Group Toolkit for Elastic Networks (NRGTEN) [17]. NRGTEN was used to calculate entropic dynamic signatures (EDS), which are vectors of predicted structural fluctuations per residue, for the docking complexes of the *µ*-opioid receptor with ligands of known efficacy generated by FlexAID. Subsequently, these EDS were used to train a LASSO linear regression-based efficacy predictor, which demonstrated strong predictive power and identified residues important for receptor activation [18].

In this work we introduce **NRGSuite-Qt**, a PyMOL plugin inspired by the first version of NRGSuite [8]. The NRGSuite-Qt, like its predecessor that is no longer functional, gives access to GetCleft, a method to detect and refine cavities in macromolecular structures, and to FlexAID a molecular docking software [5]. Additionally, NRGSuite-Qt also gives access to NRGRank [15] virtual screening simulations, the analysis of ligand-protein and protein-protein interactions with Surfaces [14], NRGTEN [17] for the calculation of dynamical signatures, generation of conformational ensembles and the effect of mutations on dynamics and stability as well as IsoMIF [12] for the detection of binding-site similarities. We also perform a comprehensive evaluation of NRGTEN/ENCoM with the SYBYL-based atom types used in FlexAID and generate distributions of Tanimoto scores of binding-site similarities to aid in the interpretation of IsoMIF results.

## 2 OVERVIEW

NRGSuite-Qt provides a user-friendly interface for the Najmanovich Research Group’s software: GetCleft, FlexAID, NRGRank, Surfaces, NRGTEN, IsoMIF, and an optional Single Mutations functionality. The descriptions of each software, their original references, and applications are listed in Table 1. NRGSuite-Qt was designed to support the use of each software individually or in combination, allowing outputs from one software to be used as inputs for another, enabling users to explore multiple workflows.

**Table 1.** Overview of tools incorporated in the NRGSuite-Qt plugin.

|  | Tool name | Functionality | Ref. |
| --- | --- | --- | --- |
| ① | **GetCleft** | Finds cavities (clefts) in protein structures. It can define specific clefts around ligands/residues, or multiple clefts in the structure. Different clefts can be defined using variations of the cleft detection parameters. Defined clefts can be refined interactively within PyMOL. It is commonly used to define binding sites for FlexAID, NRGRank, and ISOMIF. | [5, 8] |
| ② | **NRGRank** | Performs ultra-massive high-throughput screening simulations. It supports screening across predefined datasets, including FDA-approved drugs (2,500 ligands), the Chemical Component Dictionary (33,000 ligands), and all-tetrapeptides (16,000 ligands). It also allows the user to upload a SMILES ligand file. NRGRank can screen around 50 thousand ligands per-hour (8 cores). | [15] |
| ③ | **FlexAID** | A molecular docking tool for flexible ligand docking into protein targets, accounting for solvent interactions implicitly. It provides realistic docking poses and is commonly used in combination with NRG TEN to dock ligands on individual structures within conformational ensembles and Surfaces for ligand-protein interaction analysis. | [5, 8] |
| ④ | **Surfaces** | Computes pairwise molecular interaction analysis based on the calculation of surface areas in contact. This tool can be applied to *protein-ligand* and *protein-protein* interactions studies of crystal structures, computational models and docking complexes. It is commonly used for the analysis of FlexAID results and Single Mutants. | [14] |
| ⑤ | **NRG TEN** | Uses Normal Mode Analysis for the study of dynamics of proteins and protein-ligand complexes. It can calculate dynamical signatures of the system giving an overview of the flexibility in each residue. These signatures can be employed to study the effect of ligand-binding and mutations on the overall flexibility of proteins and can be used in combination with FlexAID and Single Mutations tool. | [17] |
| ⑥ | **Single Mutations** | Optional feature on NRG Suite-Qt to model single mutations using Modeller upon licensing. These mutants can be used as input for Surfaces and NRG TEN. | [21] |
| ⑦ | **IsoMIF** | Identifies molecular interaction field similarities between protein binding sites defined with GetCleft. It detects both geometric and chemical equivalences across cavities using six chemical probes. This tool can be used to predict protein function, detect potential off-targets, suggest bioisosteric replacements, and in drug repurposing efforts. | [12, 25] |

By default, three pre-processed ligand datasets are provided: CCD [11], all FDA-approved drugs available in DrugBank [19], and all tetra-peptides made by Prasasty *et al* 2019. [20]. Each software is designed to use structures loaded in the PyMOL interface, and all resulting structures are automatically loaded into the PyMOL interface to facilitate software integration. Optionally, we provide functionality for creating single mutants using Modeller [21], based on their mutate model script [22]. This functionality can be accessed through an Anaconda [23] installation, provided that the necessary licensing requirements are met.

The version of ENCoM available in NRGSuite-Qt as NRGTEN was implemented using the 40 atom-types and the respective pairwise pseudo-energy interaction matrix of FlexAID, instead of the 8 atom types and respective interaction matrix used in the original ENCoM implementation [16]. This new implementation was partially validated in the work of *Galdino et al*. [18] for the prediction of ligand efficacy in the *µ*-opioid receptor. However, that validation used a different subset of benchmarks from those used for the original ENCoM implementation as well as those from the work of *Mailhot et al*.[24] to simulate RNA dynamics. Therefore, we performed a complete analysis of the performance of this new version of ENCoM over all previously developed benchmarks.

To aid in the interpretation of IsoMIF results, we calculated the distribution of Tanimoto coefficients of binding-site similarity for all combinations of the 102 targets from DUD-E dataset [26]. We used this as a reference to calculate the Z score and the p-value for the IsoMIF results, similarly to its implementation in the Ebola VP35 study [13]. This allows users of IsoMIF within the NRGSuite-Qt to determine the statistical significance of the detected binding-site similarities.

## 3 BENCHMARKING ENCOM/NRGTEN WITH THE FLEXAID INTERACTION MATRIX

The possibility of substituting the original 8-atom type matrix employed in ENCoM, based on the LIGIN classes [27], was explored by *Galdino et al*. [18] and validated in the study of dynamics-based ligand efficacy prediction in the *µ*-opioid receptor. This modified version was implemented in NRGSuite-Qt, and its performance is here evaluated using benchmarks proposed in the original ENCoM paper [16], as well as RNA benchmarks from the work of *Mailhot et al*. [24].

The benchmarks from the original ENCoM paper include the correlation of dynamical signatures with experimentally measured b-factors, conformational change prediction calculated by the cumulative overlap of normal modes, conformational ensembles of NMR structures, and prediction of the effect of mutations on the ΔΔ*G* of folding in protein maturation. *Mailhot et al*. [24] extended these benchmarks to RNA, resulting in a total of 7 benchmarks. All datasets and metrics used in these analyses were constructed and described by *Mailhot* [28] and briefly described in the next subsections.

### 3.1 Benchmarks and datasets

For the protein b-factors correlation benchmark, we used a dataset of 80 high-resolution protein structures with a maximum length of 300 amino acids, selected from a non-redundant dataset curated by *Kundu et al*. [29], as utilized in the original ENCoM publication. For the RNA b-factors correlation benchmark, we employed a dataset of 38 RNA structures with a resolution of 2.5 Å or better, each up to 300 residues in length and with no missing atoms (benchmark available at [24]).

For each protein or RNA structure, two types of dynamical signatures were calculated, mean-square fluctuations (MSF) and entropic signatures (ES). ES are dependent on a thermodynamic scaling factor) The ES were calculated for 41 scaling factors ranging from *e−*5 to *e*^5^, and the correlation between the average b-factors of all atoms represented by a bead and their respective fluctuation values in the dynamic signature was calculated. The highest Pearson correlation value (among MSF and all 41 ES) was chosen as representative of the benchmark. We obtained Pearson correlation values of 0.569 for LIGIN atom types and 0.575 for SYBYL-based atom types.

For the protein overlap benchmark, 37 pairs of the same structure representing two forms (apo and holo) with no missing residues and the same length were used from the Protein Structural Change upon Ligand Binding Database (PSCDB) [30] as previously done [16]. Additionally, the RNA overlap benchmark consists of 227 pairs of structures of the same sequence representing two different conformations with a minimum RMSD of 2Å present in the PDB [17]. The cumulative overlap in 5% normal modes from one state to another was calculated in both directions for all pairs of RNA and proteins, and the mean cumulative overlap for both ENCoM versions is reported. The higher the overlap, the more precisely the normal modes calculated using one state, describe the movements required to reach the second state. We obtained an overlap of 0.791 and 0.793 for LIGIN and SYBYL-based atom types, respectively.

NMR solution experiments also provide a reliable source of dynamic information. To evaluate the capacity of ENCoM in capturing the movement reported by these experiments, *Mailhot et al*. [24] used Normalized Cumulative Overlap (NCO) between the motions apparent from solution NMR ensembles represented by non-rotational-translational correction to principal component analysis (nrt-PCA) and the 5% lowest frequency ENCoM normal modes of RNA alone or Protein/RNA complexes. These benchmarks designed by *Mailhot et al*. [24] contain 313 solution NMR resolved structural ensembles of RNA with at least two different states and 20 solution NMR ensembles of RNA/protein complexes with a maximum of 70% sequence variation, respectively. As with the protein overlap benchmark, a higher value is better. For the RNA-protein complexes test, we obtained 0.760 correlation for SYBYL-based atom types and 0.752 for LIGIN atom types. For RNA molecules alone, we obtained 0.773 for SYBYL-based atom types and 0.775 for LIGIN atom types. This is the only case where LIGIN atom types still show an advantage, although very small, perhaps reflecting the fact that the SYBYL-based atom type parameters were originally developed to study ligand-protein complexes and RNA interactions were not considered [5]. The optimization of pairwise interaction pseudo-energies in future studies may further improve performance, particularly for the analysis of complexes involving RNA and protein-protein interactions.

The original ENCoM [16] paper demonstrated that the predicted vibrational entropy of wild-type and mutants (Δ*S*_vib_) can be used as a linear approximation of the experimentally measured ΔΔ*G* of folding induced by mutations. To evaluate the performance of ENCoM in predicting the effect of mutations on the stability of proteins, the authors performed a linear fitting between Δ*S*_vib_ and the experimentally measured ΔΔ*G* of mutants from the ProTherm database [31] and reported the root mean square error (RMSE). This benchmark dataset contains 299 models of the mutations reported in the ProTherm database, generated from their structures available in the PDB using Modeller [21]. The smaller the reported RMSE, the better the performance of the method. The new vibrational entropy implemented in NRGTEN depends on a thermodynamic scaling factor. For each scaling factor listed in the b-factors benchmark (41 scaling factors ranging from *e*^*−*5^ to *e*^5^ as described above), we calculated the Δ*S*_vib_ for all mutants and then computed the RMSE against their experimentally measured ΔΔ*G*. We selected the scaling factor that produced the lowest RMSE as representative of this benchmark. We obtained an RMSE of 1.549 (scaling factor *e*^*−*0.5^) for the LIGIN atom types model and 1.527 (scaling factor *e*^*−*5^) for SYBYL-based atom types. Given the differences in the number and range of parameters for each of the two atom type schemes, it is natural to expect different thermodynamics scaling factors.

Table 2 summarizes all the results. The analysis of the benchmarks shows that utilizing the SYBYL-based atom types with ENCoM offers improvements in predicting protein dynamics, particularly for mutation stability prediction. Notably, the improvement in RNA b-factors correlation suggests that the new atom-type matrix is more effective in capturing RNA dynamics than the original matrix. The adoption of SYBYL-based atom types in NRGTEN generally enhances the accuracy of dynamical predictions across both protein and RNA structures, with no significant losses in performance. Since it has also been validated for the prediction of ligand efficacy in GPCRs by *Galdino et al*. [18], this version was implemented as the default version of ENCoM in the NRGSuite-Qt. With the adoption of the SYBYL-based atom types, FlexAID, NRGRank, ENCoM and Surfaces, all utilize the same atom-type scheme and pairwise atom type pseudo-energy scoring matrix.

**Table 2.** Comparison of metrics for LIGIN atom types, employed by ENCoM, and the SYBYL-based atom types, used for the FlexAID scoring function.

| Metric | LIGIN Atypes | SYBYL-based Atypes |
| --- | --- | --- |
| x Protein b-factors correlation | 0.569 | 0.575 |
| Protein Mutations RSME | 1.549 | 1.527 |
| RNA b-factors correlation | 0.446 | 0.456 |
| x Protein overlaps | 0.791 | 0.793 |
| RNA NCO | 0.775 | 0.773 |
| RNA-protein NCO | 0.752 | 0.760 |
| RNA overlaps | 0.734 | 0.728 |

## 4 DISTRIBUTION OF TANIMOTO SCORE OF BINDING-SITE SIMILARITIES

To determine whether two binding sites are significantly similar or different, we selected the dataset of all 102 targets from the DUD-E database [26]. For each ligand/target complex, we defined the cavity around the ligand using GetCleft to represent the binding site. This cavity is saved and the complex is then protonated using PyMOL’s “add hydrogens” functionality. We then selected all combinations of pairs of complexes and ran IsoMIF on each pair of cavities using the ligands as references. We calculate the distribution of Tanimoto scores of binding-site similarities for pairs of targets from different families as classified by *Mysinger et al*. (2012) [26]. We also calculate the distribution for pairs of targets from different families. Target families in DUD-E include Other Enzymes (36), Kinases (26), Proteases (15), Nuclear Receptors (11), GPCRs (5), Miscellaneous (5), Cytochrome P450 (2) and Ion Channels (2). Figure Figure 1 shows the distribution of Tanimoto scores of binding-site similarity for all 102 target, those in different or the same family, and only for kinases as the largest family with evolutionarily related proteins.

**Figure 1:**
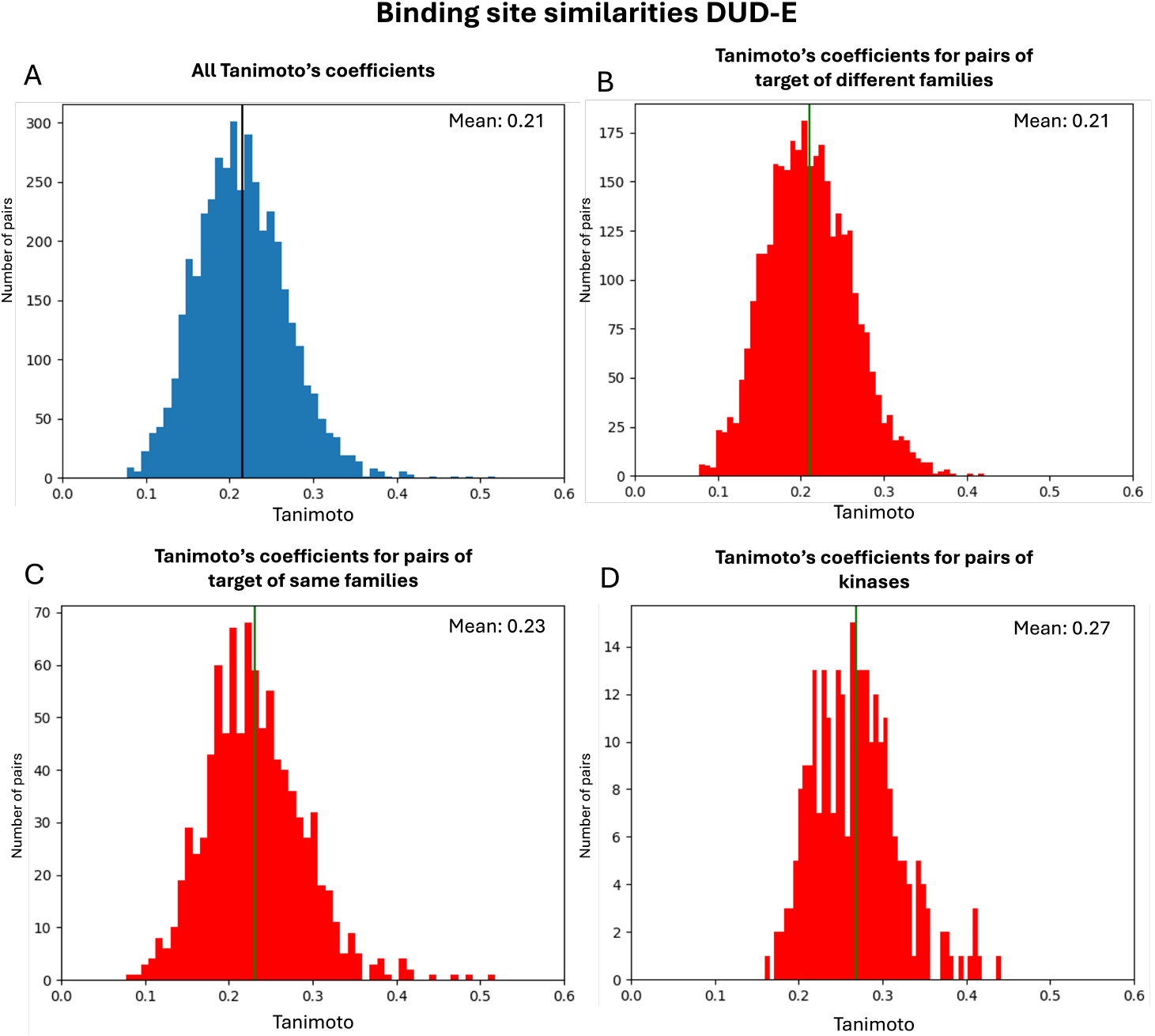
Tanimoto’s coefficient distribution for all DUD-E targets. (A) Tanimoto distribution for all pairs of targets. (B) Tanimoto distribution for target pairs of different families. (C) Tanimoto distribution for target pairs of the same families. (D) Tanimoto distribution for kinase pairs.

The mean Tanimoto coefficient for all pairs or pairs from different families is 0.21, and both distributions have a similar shape. This is not surprising as overall, most proteins in the dataset are not evolutionarily related even if belonging to the same family. For proteins of the same family, the mean is shifted to 0.23, indicating a higher similarity between proteins of the same family. The distribution is significantly shifted, with a mean value of 0.27, when considering only the kinase family, which can be explained by the fact that this family is known to have high similarity and promiscuity of ligands among different members of the family. When the user of the NRGSuite-Qt performs the detection of binding-site similarities, the Tanimoto score of binding-site similarity is plotted in relation to the distribution of Tanimoto scores across different families and serves as the basis to calculate a p-value and Z-score for the observed similarities.

## 5 CASE STUDIES

We selected two case studies (see Figure 2) demonstrating general methodologies that combine multiple tools to reproduce data previously published by *Gu S. et al*. [32] and results from two studies published by *Teruel et al*. [33, 14]. These examples were chosen to demonstrate the capability of NRGSuite-Qt to tackle complex problems such as drug repurposing, predictions of the effects of mutations on protein dynamics, and protein-protein interactions, resulting in outcomes comparable to those of the original works, but combining all analyses sequentially in a single interface. A detailed description of the case studies, including the results obtained, can be found at https://nrg-qt.readthedocs.io/. Here the focus is on presenting the general workflow for each case study.

**Figure 2:**
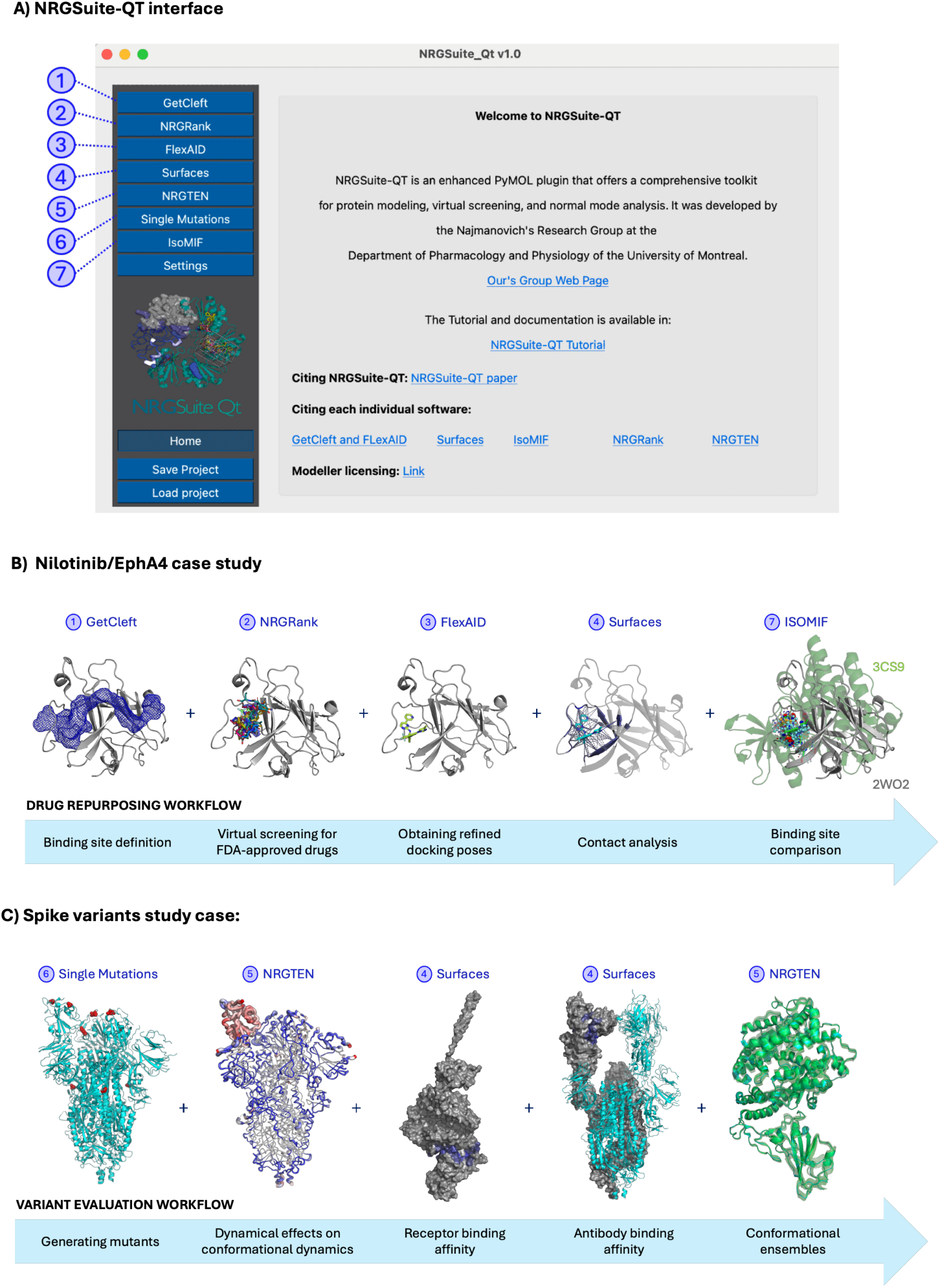
Common uses of NRGSuite. A) NRGSuite-Qt interface as a PyMOL Plugin. B) General procedure of the Nilotinib study and the workflow for drug repurposing using the FDA-approved drugs dataset in NRGSuite. C) Workflow illustrating the study of SARS-CoV-2 Spike variants and their diverse functional effects.

### 5.1 Drug Repurposing: EphA4

*Gu S. et al*. (2018) [32] screened a dataset of FDA-approved drugs for inhibitors of the receptor tyrosine kinase Erythropoietin-producing Hepatocellular A4 (EphA4), a molecular target in Alzheimer’s disease (AD). They selected and tested 22 molecules, identifying 5 potential EphA4 inhibitors. Notably, Nilotinib, a known kinase inhibitor, was found to inhibit EphA4-ephrin-A binding at a micromolar scale in a dose-dependent manner. In the NRGSuite-Qt tutorial, we used an EphA4 structure (PDB code: 2WO2), the same one used by *Gu S. et al*. in their study. First, we applied 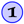 GetCleft to identify all cavities in the structure and selected the largest one by volume. We then ran 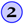 NRGRank using its predefined dataset of FDA-approved drugs. Nilotinib appeared among the top 20 results, ranked 13th, with a docking score (CF) of *−*175.54 *×* 10^4^. Next, we used 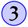 FlexAID to obtain an accurate docking pose for the Nilotinib/EphA4 complex, followed by 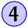 Surfaces on the top result to identify key interactions between Nilotinib and EphA4. All interactions cited by *Gu S. et al*.—including Q70, T104, I59, F154, V157, M164, L166, A193, and V159—were also observed in our docking results for Nilotinib, showing high similarity in docking poses. We can then apply 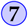 IsoMIF to calculate the similarity between the chemical environments surrounding Nilotinib in the EphA4 binding site and those of Nilotinib in complex with two other kinases: Human Mitogen-Activated Protein Kinase 11 (PDB code: 3GP0) and Human ABL Kinase (PDB code: 3CS9), both showing a Tanimoto similarity of 0.25 and 0.27 (z-score:0.78 and 1.18, respectively). This similarity is higher than the mean similarity between two random binding sites, but expected when comparing two kinases, a family for which the mean similarity is 0.27 (Figure 1).

### 5.2 Variant evaluation: Spike protein

The SARS-CoV-2 pandemic represented a pivotal moment for global scientific collaboration, driving unprecedented efforts to study the virus and its variants. Among the viral components, the Spike glycoprotein emerged as a key focus due to its high propensity for mutation, with these mutations exerting diverse functional effects. NRGSuite allows us to reproduce results from *Teruel et al*. [33, 14] to evaluate the effects of mutations on Spike’s conformational dynamics, interaction with the receptor angiotensin-converting enzyme 2 (ACE2), as well as immune recognition. Specifically, after employing the 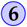 Single Mutations implementation of Modeller [21], 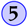 NRGTEN was applied to evaluate the effects of the D614G mutation, revealing increased flexibility in the closed conformation (PDB code: 6VXX) and rigidity in the open conformation (PDB code: 6VYB), indicating a possible contribution to the occupancy of the open conformation of Spike, needed for receptor binding and cell fusion. Mutations N501Y and K417N exhibited even greater flexibility changes in both states, which could be associated to their selection in early variants. Additionally, 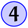 Surfaces was employed to analyze per-residue protein-protein interactions between Spike and ACE2 (PDB code: 6M17) or antibodies, such as the C105 antibody (PDB code: 6XCN). Functionally, N501Y enhanced ACE2 binding, while K417N reduced interactions with C105, potentially diminishing immune recognition. By incorporating protein ensembles generated with 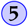 NRGTEN, we capture conformational variability to provide a dynamic and comprehensive evaluation of these interactions, acknowledging the inherent flexibility of protein structures.

## 6 CONCLUSION

NRGSuite-Qt represents a significant step forward in accessibility for structural evaluations. This accessibility comes from two key factors: the coarse-grained nature of the tools and the intuitive, user-friendly interface. The methods NRGSuite-Qt offers are inherently coarse-grained and therefore reduce the computational infrastructure required, making it ideal for high-throughput applications — a necessity as the volume of protein structure and small-molecule data continues to grow. Additionally, the user-friendly interface lowers the barrier of expertise, enabling researchers from diverse backgrounds to leverage these tools effectively.

Another pivotal feature of NRGSuite-Qt is the seamless integration of multiple tools within a single platform. This integration allows users to address complex scientific questions, as demonstrated in the case studies presented. By combining capabilities such as docking, binding site evaluation, and normal mode analysis, NRGSuite-Qt provides a cohesive environment for comprehensive structural evaluations. These features ensure that NRGSuite-Qt is not only a powerful toolkit but also a catalyst for advancing research in protein engineering and drug discovery.

## 7 AVAILABILITY

The NRGSuite-Qt is available for MacOS/Linux as well as Windows platforms. A complete installation guide and a tutorial featuring different case studies including the two shown here is available at: https://nrg-qt.readthedocs.io/

## 8 ACKNOWLEDGEMENTS

We acknowledge Olivier Mailhot for sharing his curated datasets and code for the benchmarks. We also thank the Digital Research Alliance of Canada for providing computational resources. RJN is a member of the Quebec Network for Research on Protein Function, Engineering, and Applications (PROTEO).

